# Functional Lipid Analysis via Index-Based Lipidomics Profile: A New Computational Module in LipidOne

**DOI:** 10.1101/2025.11.04.686489

**Authors:** H. B. R. Alabed, D. F. Mancini, M. Pergola, L. Romani, S. Martino, A. Koulman, R. M. Pellegrino

## Abstract

Understanding the functional roles of lipids is essential for interpreting metabolic phenotypes in health, disease, and dietary interventions. Here, we present a major update to LipidOne, a user-friendly web-based platform for lipidomic data interpretation (lipidone.eu), introducing the novel analytical module: Functional Lipid Analysis (FLA).

This component enables the assessment of the quantitative features of lipidomic datasets through a biologically structured, index-based approach. The FLA module computes 42 indices representing specific lipid function— including membrane structure, energy storage, and signaling. These indices are derived from lipid classes, molecular species, and fatty acyl-, alkyl-, and alkenyl-chain composition. Each index is statistically compared across experimental groups and analyzed and visualized through dedicated tools, including bar plots, volcano plots, PCA, PLS-DA, heatmaps, and functional radar charts.

Every index is semantically annotated with biologically meaningful phrases, allowing users to move beyond numerical variations and toward mechanistic insights of lipid function. In the FLA module, index variations are further linked to predicted protein mediators, bridging lipid alterations to enzymatic pathways and enabling network-based interpretations. This integrative strategy lays the foundation for a systems biology perspective, connecting lipidomics to proteomics and or transcriptomics yielding functional pathway analysis.

We demonstrate the utility of this framework using datasets from two published works. In both cases, FLA confirmed the authors’ conclusions and yielded additional, biologically coherent functional readouts not originally emphasized.

By shifting the focus from individual lipid species to interpretable biochemical indices, LipidOne 2.3 offers a reproducible, scalable, and biologically informed platform for systems-level lipid biology and hypothesis generation.

## 1. Introduction

Lipids constitute a highly diverse group of biomolecules that are traditionally classified into their major categories based on their structure, each containing multiple classes and subclasses of structurally related species. This complexity arises from their modular architecture: lipid molecules are assembled from distinct building blocks—such as glycerol, acyl-alkyl-alkenyl-chains, sphingoid bases, phosphate groups, and saccharides—each with a specific metabolism, enzymatic regulation, and subcellular distribution. Despite their diversity, different biosynthetic routes can converge to produce identical or functionally similar lipid molecules, reflecting a high degree of redundancy and plasticity within lipid metabolic networks (Fahy *et al*., 2005).

### From Lipid Complexity to Biological Insight: A Computational Framework for Functional Lipidomics

The biological roles of lipids are equally diverse. They act as essential components of cellular membranes, which makes them essential to cellular structures as well as regulating membrane protein function, as reservoirs of metabolic energy, and as powerful signaling mediators involved in virtually every physiological process. Importantly, lipid functions can be exerted at multiple levels: entire classes (e.g., triglycerides, phospholipids), individual molecular species (e.g., PE 16:0/22:6), or even specific substructures such as fatty acyl chains or headgroups may carry distinct and sometimes independent biochemical implications (Conroy *et al*., 2024).

Lipids are highly dynamic molecules that undergo continuous remodeling in response to environmental, physiological and pathological stimuli. Compared with aqueous (polar) metabolites, which can fluctuate rapidly and transiently, lipid species generally change on slower timescales when considering membrane composition and neutral-lipid storage/usage. Nevertheless, specific lipid signaling steps can be fast (e.g., COX- and LOX-mediated eicosanoid production) and also the localization of lipid in membranes can be highly dynamic. Because lipids frequently sit at entry and exit nodes of complex metabolic networks, they integrate upstream and downstream biochemical signals; the cumulative effects of these changes are particularly relevant to chronic diseases and long-term adaptation to physiological stress (Cho *et al*., 2023).

Moreover, lipid biochemistry is tightly interwoven with protein signaling pathways and the metabolism of water-soluble metabolites, forming a complex regulatory network that governs cell fate decisions, membrane organization, and energy homeostasis (Sych *et al*., 2022).

Over the past decade, lipidomics has emerged as a powerful omics approach to quantitatively map the lipid composition of biological systems. Typically, a lipidomic study focuses on one or more of the following applications:

- Biomarker discovery using statistical and bioinformatic analysis, to identify lipid-based markers for early diagnosis, disease progression, or therapeutic response—especially valuable in cardiometabolic and neurodegenerative disorders (Hornemann, 2022).
- Pathway analysis, which seeks to reconstruct lipid transformation routes and infer the involvement of specific genes, enzymes, and regulatory proteins by integrating lipidomic profiles with transcriptomic or proteomic data (Nguyen *et al*., 2017).
- Functional analysis, which focuses on interpreting the biological roles of lipids based on their variation across conditions, and on linking those changes to cellular behaviors such as membrane remodeling, energy shifts, or inflammatory signaling.

However, unlike biomarker discovery and pathway reconstruction, functional interpretation remains the main bottleneck in the utilization of lipidomics. The enormous diversity of lipid species and the limited coverage of curated knowledge bases (KEGG, Reactome, WikiPathways, HMDB) leave many measured species unmapped, hampering downstream functional inference. Existing platforms such as LipidOne (8) and BioPAN (9) mitigate pathway gaps. Here we address the unmet need: FLA translates quantitative lipid changes into a concise set of biologically grounded indices that capture structural, energetic, and signaling functions across conditions.

When it comes to functional analysis, it is essential to recognize that lipids perform three fundamental biochemical functions in living organisms: structural, energetic, and signaling/regulation (Uzman, 2001). This triad is a foundational concept in lipid biochemistry, but it has not yet been fully translated into computational tools capable of resolving which of these functions are altered in a given experimental context (Shevchenko and Simons, 2010). A few platforms, such as LION/web (Molenaar *et al*., 2019), MetaboAnalyst (Lu *et al*., 2023) and LipidSig (Lin *et al*., 2021), are excellent resources that offer, among other features, tools described as “functional analysis” for lipidomic datasets. However, in all these contexts, the term refers primarily to enrichment-based approaches, conceptually derived from transcriptomics and proteomics, and does not entail direct biochemical inference—such as distinguishing the structural, energetic, or signaling roles of lipids within specific experimental conditions. As a result, none of these platforms provides a mechanistic or context-aware interpretation of lipid function, revealing a critical gap in the current bioinformatics landscape for lipidomics.

This gap is especially relevant because lipid metabolites, due to their chemical heterogeneity, functional redundancy, and compartmental specificity, require dedicated analysis strategies. Interpreting lipidomic changes purely in terms of class enrichment or fold changes is often insufficient and may lead to misleading conclusions. This is particularly true for functional analyses that adopt enrichment-based strategies originally developed for gene expression data. As highlighted by Lee et al., such approaches can misrepresent the biology of small molecules like lipids, which differ substantially from genes in structure, dynamics, and regulation (Lee *et al*., 2025). What is needed is a framework that integrates quantitative lipid features, biochemical functions, and systems-level reasoning to generate interpretable biological outputs.

To address this unmet need, we developed a dedicated module within the LipidOne platform (Pellegrino *et al*., 2022; Alabed *et al*., 2024) that systematically translates quantitative lipidomic data into biochemically interpretable functional profiles. The Functional Lipid Analysis (FLA) module provides a structured and reproducible approach to identify whether specific lipid functions are enhanced or suppressed across experimental conditions.

## 2. Methods

### Functional Lipid Analysis (FLA) module

The Functional Lipid Analysis (FLA) module within LipidOne is based on a curated set of 42 mathematically defined indices, which are quantified from user-uploaded lipidomics datasets. These indices are calculated as ratios between lipid species, subclasses, or defined building blocks—including acyl, alkyl, or alkenyl chains, and headgroups. Each index is specifically designed to capture distinct biochemical alterations in lipid function, reflecting properties such as the saturated/unsaturated lipid ratio, triglyceride metabolism, or the relative abundance of signaling lipids like ceramides.

To facilitate interpretation, the indices are grouped into three major functional categories, corresponding to the principal roles of lipids in biological systems (Gurr *et al*., 2016):

- **Structural indices**, which describe both quantitative and qualitative features of membrane-associated lipids. These indices capture variations in the abundance of entire lipid classes (e.g., phosphatidylcholines, cardiolipins), but also reflect molecular-level properties such as the degree of unsaturation (linked to membrane fluidity), average chain length (affecting membrane curvature), and the presence of ether bonds. While certain structural lipids like cardiolipins are localized in mitochondria and have multifunctional roles, the structural indices in this group aim to represent the overarching biophysical properties of lipid assemblies across different cellular compartments.
- **Signaling indices**, which track lipid species and subclasses involved in cell signaling, inflammation, and intercellular communication. These include not only individual bioactive molecules—such as omega-3 and omega-6 polyunsaturated fatty acids and eicosanoid precursors—but also broader lipid classes or sub-classes known for their signaling functions, including ceramides, lysolipids, and ether lipids. These molecules act as metabolic messengers, modulating immune responses, stress signaling, and pathways related to cell growth, differentiation, or apoptosis.
- **Energy-related indices**, which capture lipid species involved in metabolic energy storage, mobilization, and mitochondrial function. These include not only neutral lipids such as triglycerides and diglycerides, but also lipid classes and acyl chain features linked to β-oxidation. For instance, changes in acyl chain length may reflect enhanced or impaired fatty acid catabolism, while specific mitochondrial lipids—such as cardiolipins and acyl-carnitines—are functionally associated with energy production and transport (Grevengoed et al., 2014; Paradies et al., 2019). Although some of these, like cardiolipins, may also contribute to structural integrity, their metabolic implications justify their partial inclusion in this group.

See Table S1 for more information.

Each index in Table 1 has been systematically curated based on evidence from the scientific literature and reflects established relationships between lipid composition and cellular functions. Indices are also linked to specific enzymes or regulatory proteins that determine, modulate or respond to lipid variation. For instance, the Ceramide/Sphingomyelin Index is associated with enzymes involved in apoptotic signaling and stress responses such CerS1, CerS2, SMS1, SMS2, SPTLC1, while the DG/TG Index reflects lipid mobilization and energy storage dynamics by activation of ACSL1, DGAT2, LPL, AGPAT2, LCAT proteins. By linking these indices with the associated proteins, LipidOne the opportunity to make inferences about the mechanisms that drive the differences between the analyzed samples.

**Table 1:**
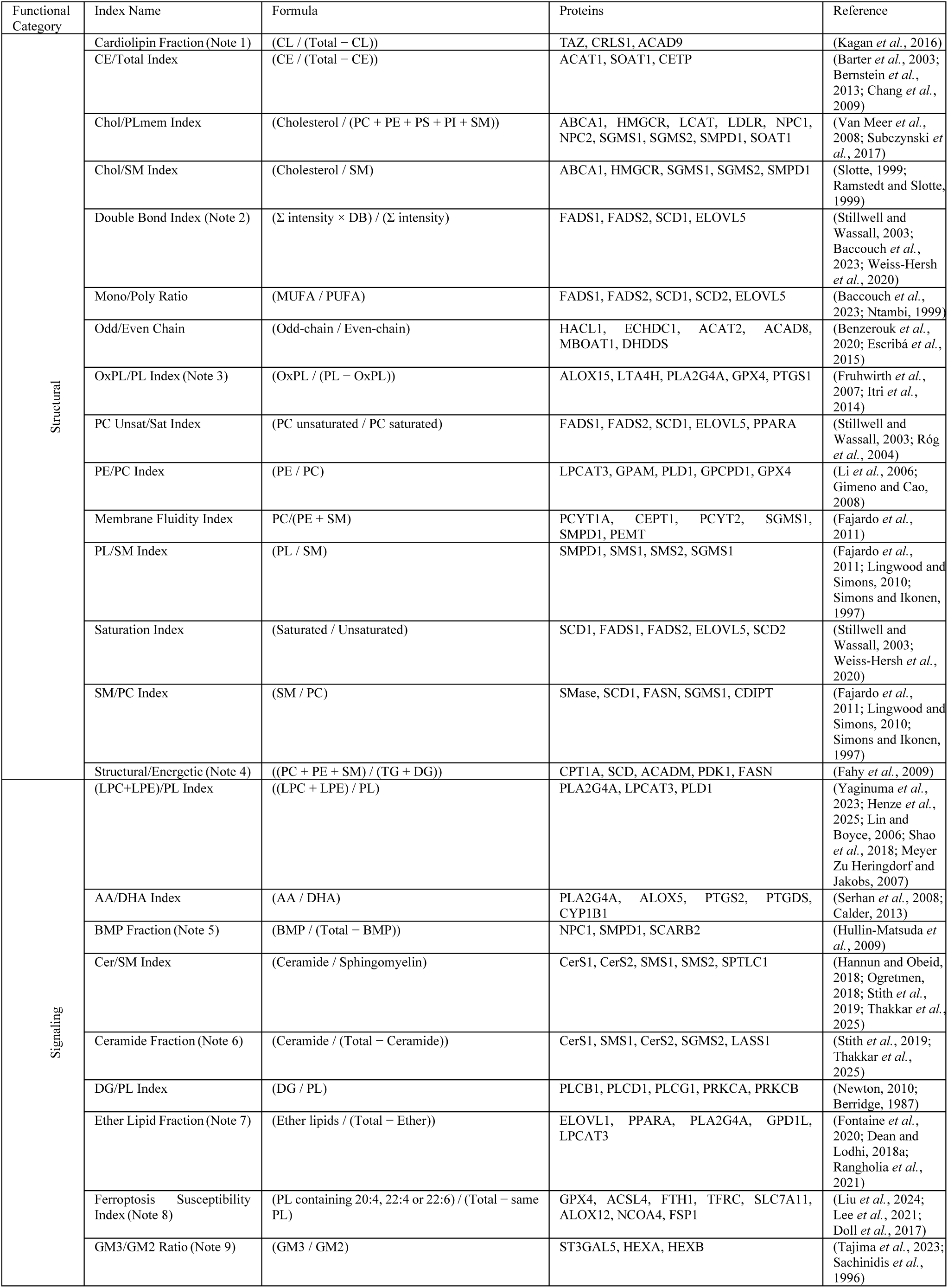
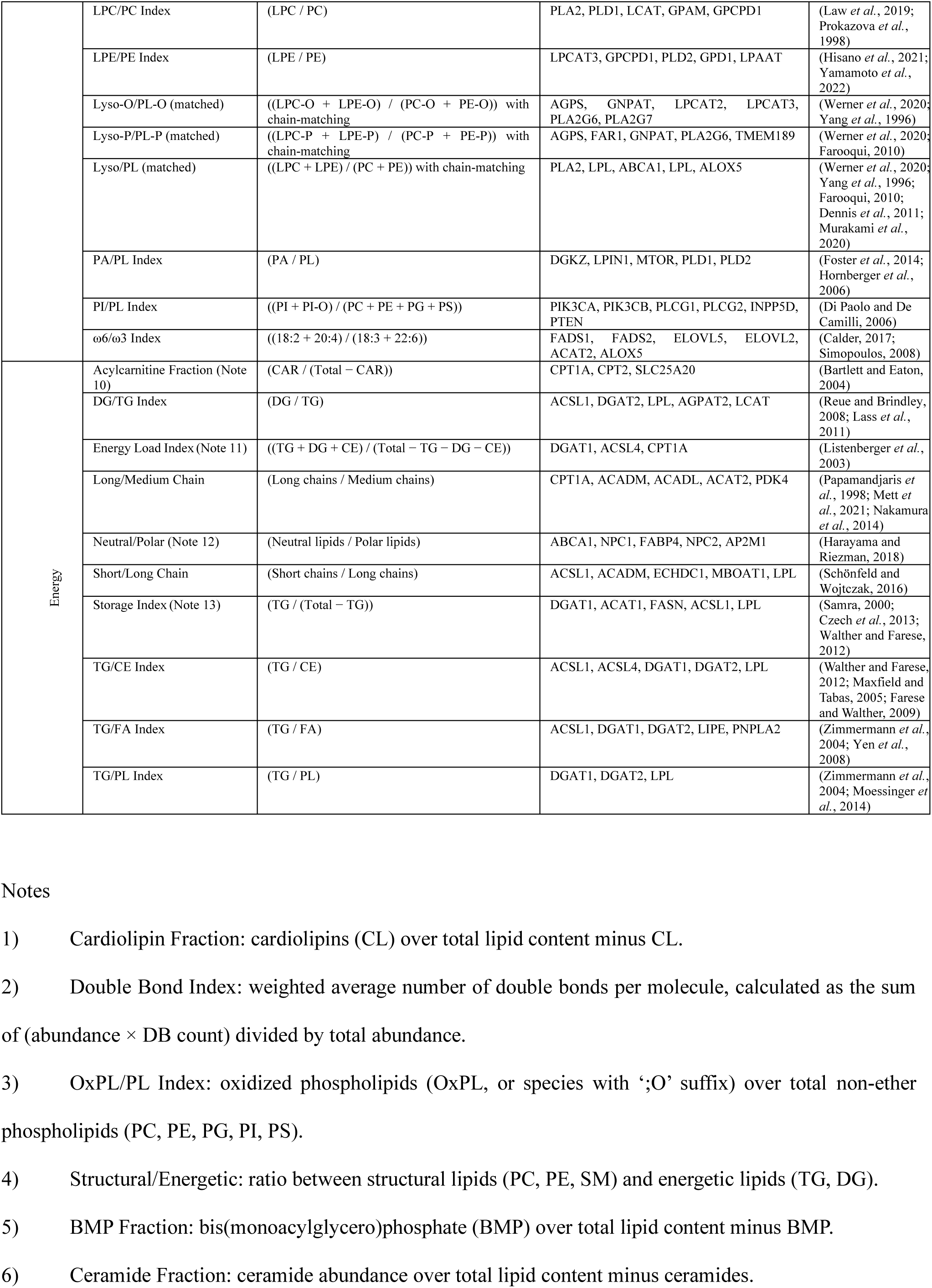
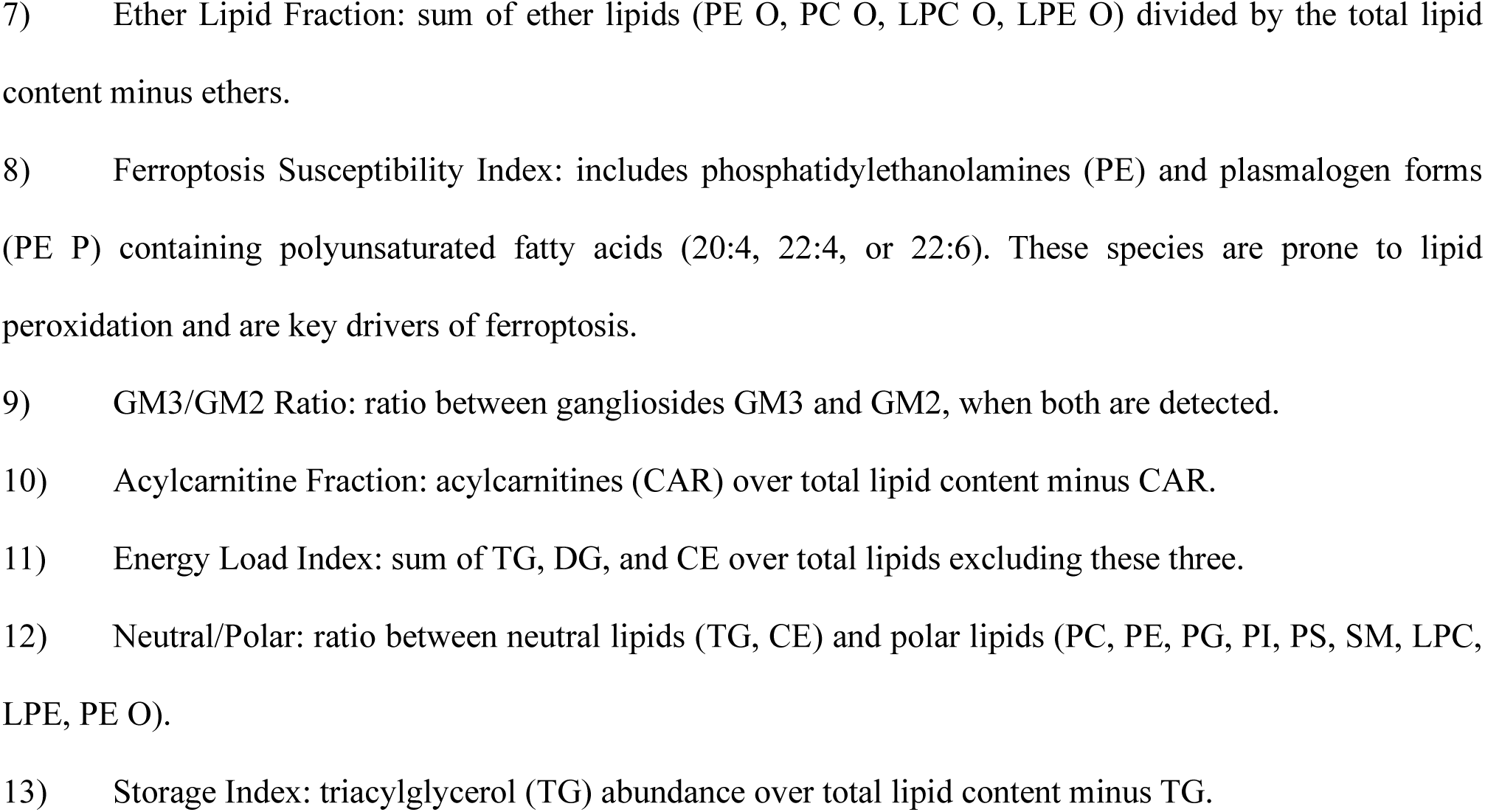
Functional lipid indices, biochemical classification, and calculation formula

As the indices are directly related to specific proteins the FLA module facilitates the construction of protein interaction networks, based on predicted activations or deactivations of these proteins, allowing researchers to visualize how specific lipid alterations might influence broader biological pathways. The network-based interpretation extends to protein clusters, where specific protein families—such as kinases or phosphatases— may be identified as central mediators of lipid-driven pathways. By visualizing these interactions, users can generate hypotheses regarding the functional consequences of lipidomic changes. These insights can then inform experimental designs or suggest potential therapeutic interventions.

Finally, since LipidOne has been designed to build protein interaction networks for 10 model organisms, we provide homologous proteins from these species based on their short names in the STRING database. These are listed in Table S2.

The full set of 42 curated functional indices is automatically computed for each sample in the lipidomics dataset when the user accesses the Functional Lipid Analysis (FLA) module of LipidOne. The use of ratio-based definitions ensures that the indices remain interpretable, scalable, and robust across experimental platforms, minimizing dependency on absolute quantification. The resulting numerical values serve as variables for statistical analysis and group comparison enabling the identification of biologically meaningful differences across experimental conditions, as illustrated in the following sections.

### FLA workflow

The FLA module integrates statistical and visualization tools to detect and interpret alterations in lipid function across experimental conditions. All analyses are implemented in R and executed server-side, ensuring reproducibility, transparency, and scalability. Starting from the user’s lipidomics dataset, the platform computes a panel of 42 mathematically defined functional indices, which act as the primary variables. These indices are statistically compared across groups (typically with t-tests or ANOVA), and the resulting log₂ fold changes and p-values drive the plots and summaries. The application on these analyses will be shown further down in the text.

As the general philosophy of LipidOne, the tools of FLA module, have been grouped into four categories:

## 1. Univariate Statistical Analysis

- **Functional Summary:** Compares all indices between two selected groups. **Bar Plot** show the group mean log2 fold change, while the error bars depict the standard error of the log2 fold change estimated by first-order error propagation (delta method) from within-group variability and sample size. This Standard Error-based approach is coherent with the Welch test used for significance and provides a compact, distribution-light estimate of uncertainty. Bars are colored according to the predominant lipid function of each index (Figure S1.A). **Radar Plot** show a companion radar chart summarizes the overall functional landscape (Figure S1.B). **Functional Dominance** show a three-bar panel (Structural, Signaling, Energy) that reports, at a glance, the prevailing functional shift (Figure S1.C). For each function, we aggregate the magnitude of change by summing the absolute log2FC of indices that pass significance and normalizing by the number of indices effectively evaluated in that function, to avoid bias from missing/filtered indices. Larger bars indicate a stronger net deviation of that function in the experimental group. Functional Summary outputs are accompanied by CSV exports reporting per-index statistics (log2FC, SE, p-value from Welch’s test, and other metrics when available). For indices that meet significance thresholds, the CSVs also include a standardized Interpretation field with concise phrases tailored to the index identity and direction of change, enabling immediate reuse in figure tables and narrative summaries.
- **Functional Volcano Plot**: Highlights the magnitude of change (log₂ fold change) and significance for each index. In FLA, a three-panel volcano separates indices by functional category. A results table lists indices that are both significant and beyond the user-defined variation threshold (Figure S1.D). The significant indices are presented together with brief sentences explaining the biochemical meaning of the differences.
- **Functional Bubble Plot:** Displays a bubble heatmap of the top ten indices across groups; bubble size reflects significance and color encodes fold change. Each index is accompanied by a concise interpretive sentence (Figure S 1.E).
- **Functional Biomarker Discovery**: Ranks indices by discriminative performance (p-value, ROC AUC— polarity-corrected when needed—statistical power, Cohen’s d). Outputs a Top-20 table (CSV + PNG) for rapid screening and prioritization (Figure S1.F).
- **Functional Correlation Circoplot**: Visualizes pairwise correlations between indices to reveal co-regulation and functional synergy (Figure S1.G).

## 2. Multivariate Statistical Analysis

Instead of analyzing hundreds or thousands of molecular lipid species, FLA works with a compact set of functional indices (currently 42) that explicitly encode lipid roles (structural, energetic, signaling). This biochemistry-driven reduction in data complexity makes principal components and latent variables easier to interpret, because each feature reflects a function rather than an individual species and reduces co-linearity that often occurs in lipidomics data. By aggregating related species, the indices improve signal-to-noise and often sharpen group separation in PCA and PLS-DA. (see details in in the results section). Finally, VIP scores and loadings calculated on indices map directly onto functional biochemical mechanisms, revealing unexpected links between lipid functions.

- **Functional PCA:** PCA on the matrix of functional indices (centered and auto-scaled by default; scaling options are user-configurable). Produces score plots (sample clustering) and loading plots (index contributions), enabling dimensionality reduction and identification of dominant functional axes (Figure S1.H).
- **Functional PLS-DA:** Performs Partial Least Squares Discriminant Analysis on the functional index matrix.

Outputs include score plots with confidence ellipses (Figure S1.I), loading plot (Figure S1.J), and VIP score bar plots (Figure S1.K). This supervised method improves discrimination between experimental groups and highlights key functional biomarkers.

## 3. Clustering Analysis

• **Functional HeatMap:** Creates a clustered heatmap of indices across all samples of groups selected (Figure S1.L). Hierarchical clustering is applied to both samples and indices. User can select the number of indices to include and apply one of seven diverse algorithms of distances, aiding in the identification of group-specific functional or nutritional profiles and outlier patterns.

## 4. Lipid System Biology

• **Predicted Functional Proteins Network:** This tool links each lipid functional index to relevant proteins and enzymes using the STRING database (Figure S1.M). The resulting network graph highlights biological pathways and regulatory modules, with nodes colored according to cluster annotations, thus providing a mechanistic link between lipidomic alterations and the proteins potentially involved in their regulation or response. To enhance interpretability and flexibility, the user can adjust key parameters of the analysis, including the statistical thresholds used to define significant indices (p-value and log₂ fold change), the type of STRING interaction network to be queried (full network or physical interactions only), and the confidence score threshold for STRING interactions. Additionally, users can specify the number of neighboring proteins to retrieve for each lipid-associated hit and choose the annotation system used to color the protein nodes in the network. Available annotation sources include Gene Ontology (Biological Process, Molecular Function, and Cellular Component), KEGG Pathways, Pfam Domains, InterPro Families, UniProt Keywords, and Reactome Pathways. This high degree of configurability allows users to tailor the network reconstruction to the specific biological context of their study, supporting hypothesis generation and functional interpretation of lipidomic data.

## 3. Result and discussion

To demonstrate the power of the FLA module in detecting biologically meaningful alterations in lipid metabolism, we applied it to a lipidomic dataset from a study investigating the role of PDK1 in cardiomyocyte lipid utilization (Atser *et al*., 2025).

**Functional Lipid Analysis captures impaired lipid utilization upon Phosphoinositide-dependent kinase- 1 knockdown in cardiomyocytes**

In this study, the authors reported that Phosphoinositide-dependent kinase-1 (PDK1) knockdown (KD) led to impaired lipid mobilization and reduced energy buffering compared to scrambled controls (SC), as evidenced by changes in lipid classes and metabolic markers. They observed a shift towards enhanced lipid storage and reduced fatty acid oxidation, indicative of a metabolic imbalance.

To explore the global lipid functional landscape in response to PDK1 knockdown, we performed principal component analysis (PCA) using the set of functional lipid indices computed by the FLA module.

The score plot revealed a clear separation between KD and SC along PC1 (72.9% of variance), capturing the dominant functional difference in the dataset. The loading plot confirmed that energy-related indices—Storage Index, TG/PL Index, Energy Load Index—were the main contributors, aligning with the KD condition. In contrast, structural and signaling indices contributed more modestly along PC2, indicating that the primary alteration is related to energetic lipid turnover.

### Functional Summary and Lipid Domain Shifts

Using the Functional Summary feature, we compared all indices between KD and SC cells under nutrient deprivation. This analysis revealed a selective shift across the three major lipid domains—structural, signaling, and energy.

Notably, while structural lipids exhibited modest and heterogeneous changes, energy-related lipid indices were strongly upregulated in KD. The Storage Index, TG/PL Index, and Energy Load Index showed significantly positive log2 fold-changes (Figure 2), consistent with the experimentally observed impairment in triacylglycerol hydrolysis. By contrast, the DG/TG Index was significantly decreased, consistent with a relative accumulation of triacylglycerols over diacylglycerols and a reduced lipolytic flux. Together with the increases in the Storage Index, TG/PL Index and Energy Load Index, this pattern indicates enhanced neutral-lipid storage but a diminished capacity to mobilize stored lipids into readily oxidizable substrates (significance as indicated in Fig. 2).

**Figure 1:**
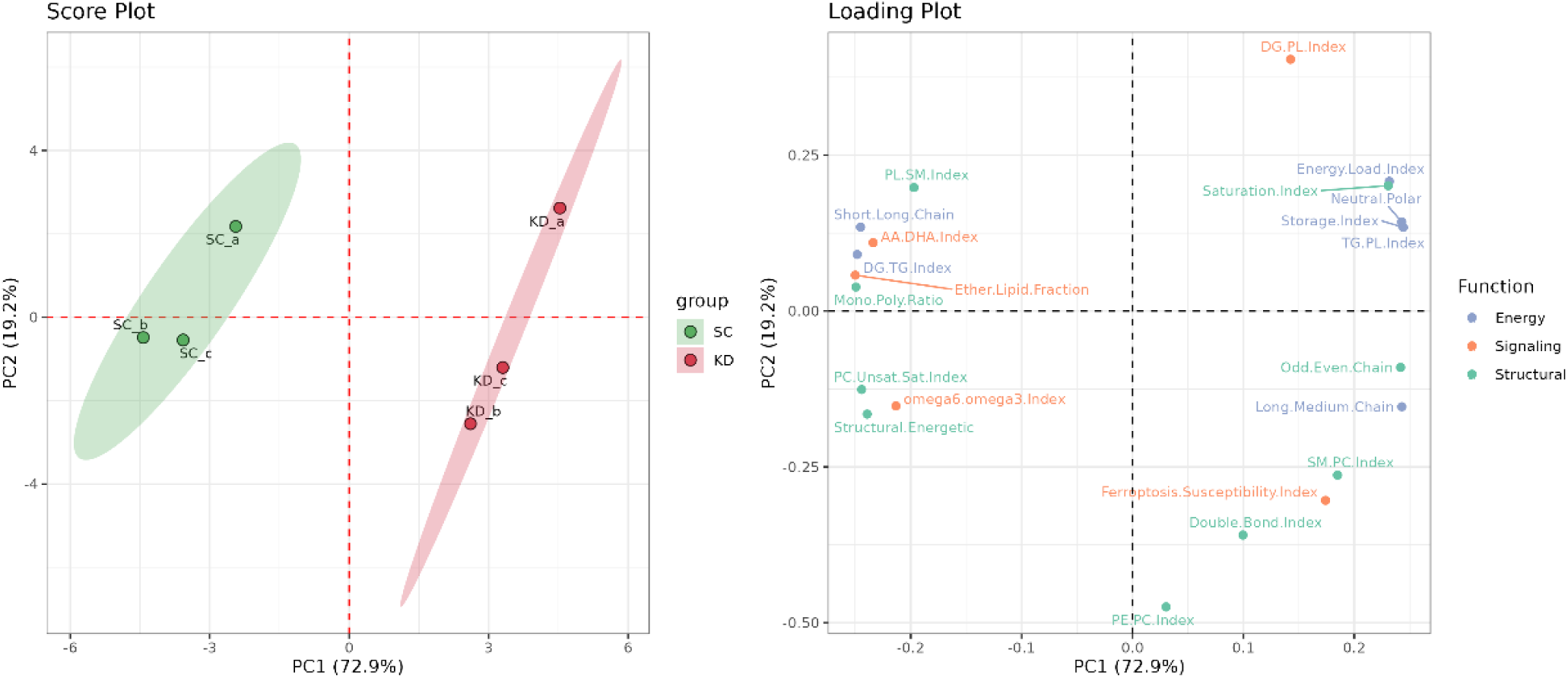
Principal component analysis of functional lipid indices in PDK1 knockdown (KD) versus control (SC) cardiomyocytes: Left: The score plot shows a complete separation between KD and SC samples along PC1 (72.9% of variance), with minimal intra-group variability. Right: The loading plot indicates that energy-related indices (e.g., Storage Index, TG/PL Index, Energy Load Index) strongly contribute to the first principal component, in agreement with the lipid functional shift observed in KD cells. Structural and signaling indices show more modest contributions along PC2 (19.2%), confirming that the dominant functional alteration lies in energy lipid metabolism.

**Figure 2:**
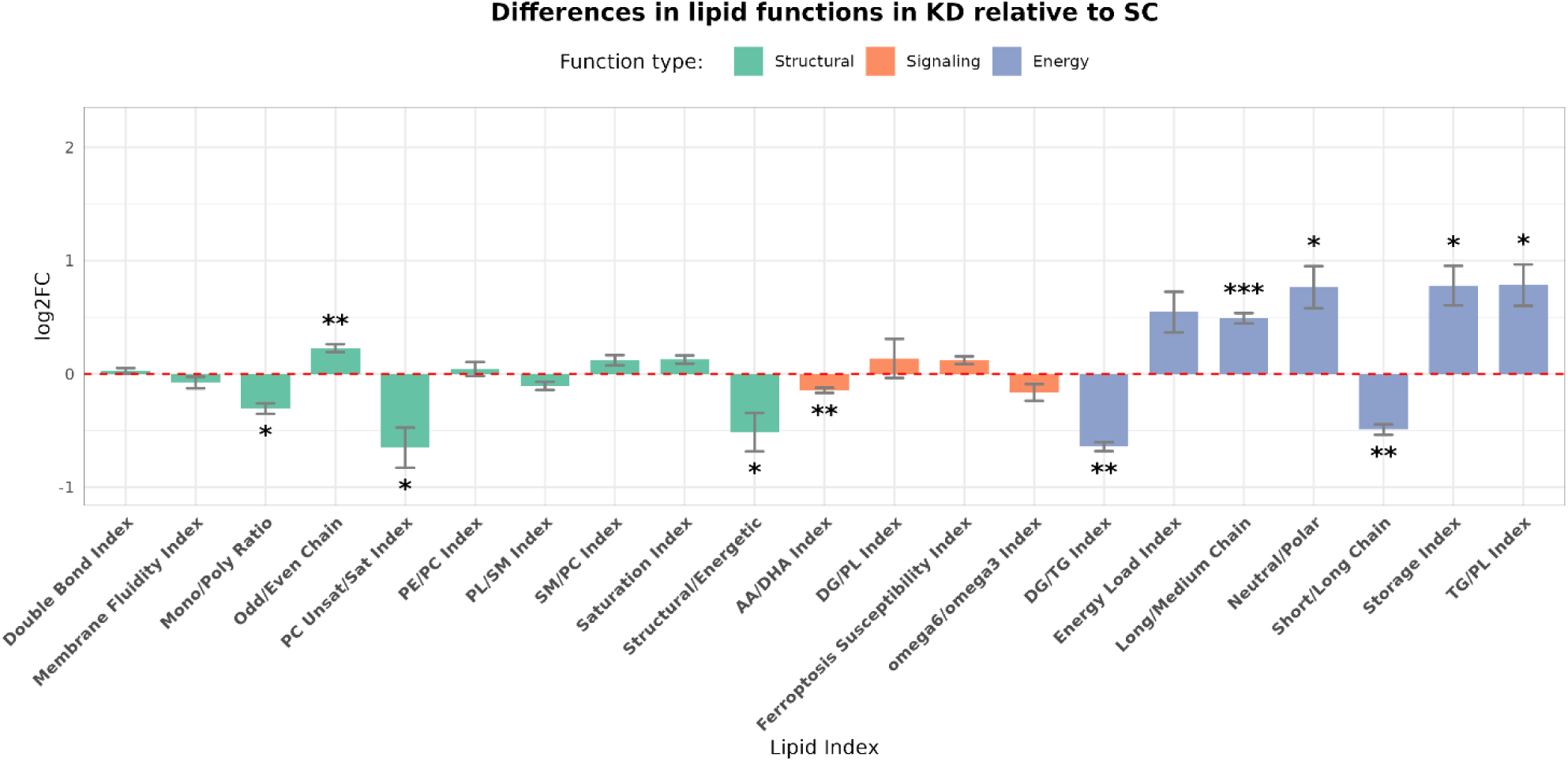
Differences in functional lipid indices in PDK1 knockdown (KD) versus scrambled control (SC) cardiomyocytes under nutrient deprivation. Bar plot showing log₂ fold-change (log₂FC) values for functional lipid indices calculated by the FLA module, grouped by lipid function: structural (green), signaling (orange), and energy (blue). Positive values indicate higher index values in KD, negative values indicate lower values relative to SC. KD cells exhibited a pronounced upregulation of storage-related energy indices (e.g., Storage Index, TG/PL Index, Long/Medium Chain), alongside significant downregulation of mobilization-related indices (Energy Index, DG/TG Index), indicative of enhanced lipid storage but reduced lipolytic flux and energy release capacity. Selected signaling indices (Ether Lipid Fraction, AA/DHA Index) were also significantly decreased, suggesting altered lipid-mediated signaling and potential anti-inflammatory adaptation. Asterisks denote statistical significance (*p < 0.05; **p < 0.01; ***p < 0.001).

**Figure 3:**
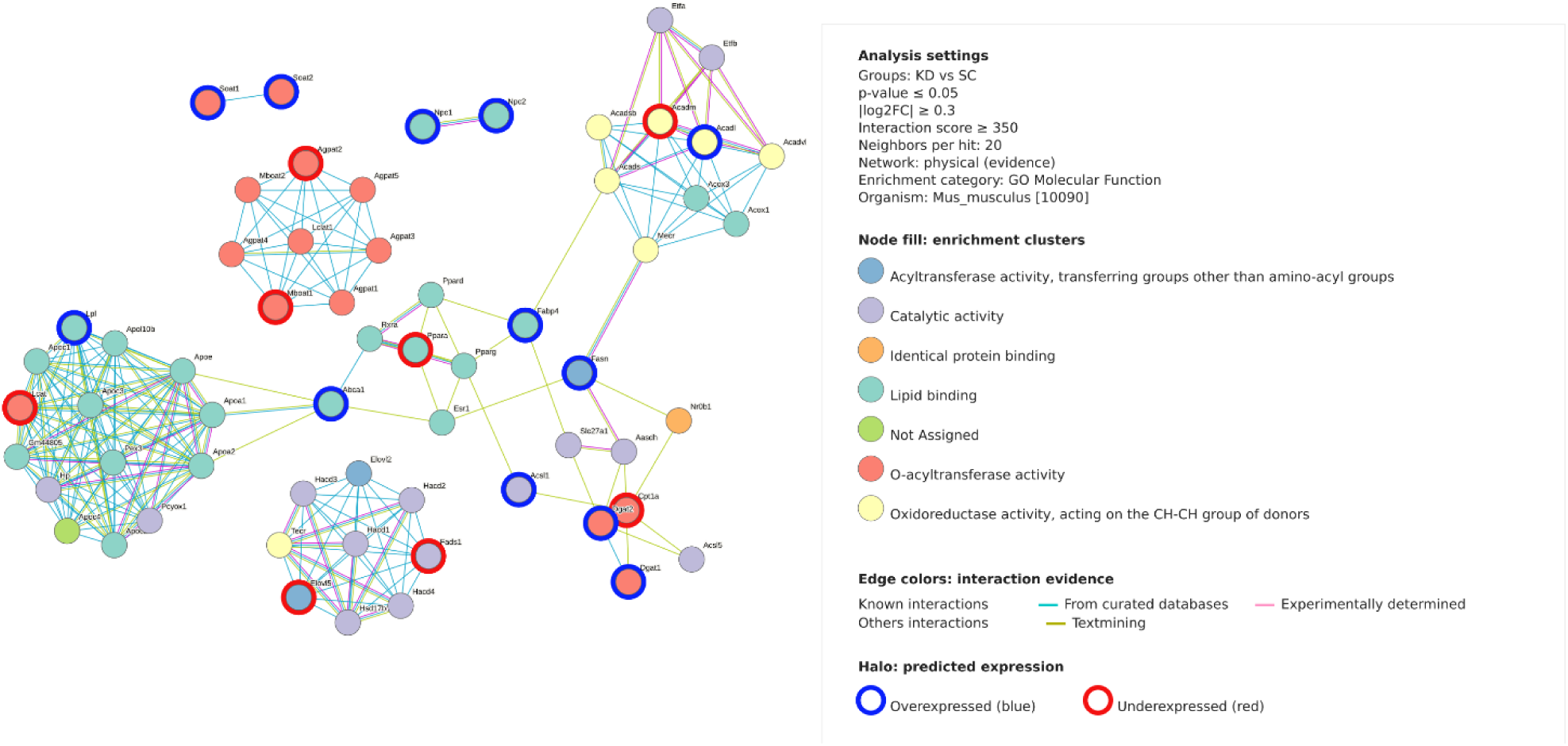
FLA-guided lipid–protein interaction network in mouse cardiomyocytes after Pdk1 knockdown. Nodes are mouse protein symbols; edges are STRING functional associations (edge color = evidence channel). Node fill = GO Molecular Function enrichment clusters. Node halos = FLA-predicted expression status (blue, overexpressed; red, under-expressed). The layout highlights four major regions: FAO/β-oxidation (left), lipid-binding/lipoprotein cluster (upper), acyl-chain editing (right, e.g., Fasn, Scd1, Elovl5/6, Fads1/2), and acyltransferases/glycerolipid remodeling (lower-right, e.g., Agpat, Lclat1, Lpcat, together with Lpin/Dgat). This organization aligns with FLA’s functional readouts indicating increased storage alongside reduced fatty-acid utilization in KD.

Interestingly, certain signaling lipid indices—such as the Ether Lipid Fraction and AA/DHA Index—were significantly downregulated. This suggests diminished ether lipid synthesis or turnover, potentially altering response to oxidative stress (Jové *et al*., 2023; Perez *et al*., 2022) and changing specific lipid-mediated cellular pathways (Dean and Lodhi, 2018b). The reduced AA/DHA ratio could suggest an alteration in anti or/and anti-inflammatory processes. Such findings highlight the added value of a function-based approach, which can uncover subtle, yet biologically relevant shifts often missed in compositional or class-based analyses.

### Molecular mechanisms and protein-lipid interaction network

To probe mechanism, we built an FLA-guided STRING network (mouse protein symbols). Node fill encodes GO Molecular Function enrichment clusters, edge colors show STRING evidence channels, and node halos report the FLA-predicted direction (blue, overexpressed; red, under-expressed). The layout reveals four coherent modules:

FAO/β-oxidation. Enzymes for activation, import and mitochondrial β-oxidation (e.g., Acsl, Cpt1/2, Acadvl/Acadm, Eci1, Decr1, Etfa/b, Hadha/b, Acaa1/2). The mixed halo pattern, with several under-expressed nodes, is consistent with the FLA inference of lower mitochondrial FA utilization in KD.

**Lipid-binding/lipoprotein cluster**. A densely connected group of lipid-binding proteins and apolipoproteins, indicating changes in lipid handling and trafficking at the interface with extracellular/particle-associated pathways.

**Acyl-chain editing**. De novo synthesis, elongation and desaturation (e.g., Fasn, Scd1, Elovl5/6, Fads1/2, Hsd17b12), consistent with adjustments in chain length and unsaturation that accompany the storage-dominant phenotype.

**Acyltransferases/glycerolipid remodelling**. Lysophospholipid and glycerolipid acyltransferases (e.g., Agpat family, Lclat1, Lpcat), together with phosphatidate/diacylglycerol branch enzymes (e.g., Lpin, Dgat), supporting enhanced neutral-lipid storage and ongoing phospholipid remodeling.

Across modules, the network maps FLA-inferred functional shifts onto enzyme-level hypotheses (e.g., the Pnpla2/ATGL axis and the FAO machinery), providing a mechanistic scaffold for follow-up experiments.

The FLA results agree with the original study showing that PDK1 knockdown impairs lipid mobilization and fatty-acid oxidation (FAO). Beyond this confirmation, FLA resolves additional features: indices indicate concurrent lipid storage increase with reduced mobilization (e.g., Energy Index, DG/TG), consistent with a lower lipolytic flux and limited rapid energy release. It also detects shifts in signaling-lipid indices and predicts coherent changes in the protein network, meriting follow-up.

In the same dataset, FLA recapitulates the drop in TAG hydrolysis, the reduced mitochondrial FAO capacity, and the predicted decrease in ATGL activity in KD cells. Moreover, it captures functional shifts that parallel the subtle remodeling of TG acyl-chain length and unsaturation reported by the authors—patterns that bulk compositional views can miss. Thus, FLA reproduces the study’s in vitro/in vivo convergence directly from lipidomics, offering a rapid, mechanism-oriented readout of the phenotype.

Finally, FLA adds mechanistic resolution by quantifying the divergence between storage-positive (Storage, TG/PL, Energy Load) and mobilization-negative (Energy, DG/TG) indices; by revealing organelle-aware changes (elevated LPE/PE consistent with inner-mitochondrial remodeling, reduced ether-lipid and AA/DHA signals); and by mapping these alterations onto a Pdk1–Pdk4–Pdha1-centred network connected to Pnpla2 (ATGL) and the FAO machinery. Together, these results show that a single lipidomics dataset can yield a function-based interpretation that both confirms impaired lipid utilization and pinpoints enzyme-level hypotheses underlying the cardiomyocyte phenotype—within minutes.

### Functional Lipid Analysis of lipidomic signatures in Hepatocellular carcinoma and chronic hepatitis C virus -related conditions

Hepatocellular carcinoma (HCC) is the most common primary liver malignancy, often arising in patients with chronic hepatitis C virus (HCV) infection. Early diagnosis remains a clinical challenge, particularly in AFP-negative patients, where traditional biomarkers and imaging approaches show limited sensitivity. In a recent multi-omics investigation, Caponigro et al. (2023) (Caponigro *et al*., 2023) performed integrated untargeted metabolomics and lipidomics profiling of plasma from 102 HCV-positive subjects, including patients with HCC (n = 69), chronic HCV infection without cancer (n = 23), and HCV-associated mixed cryoglobulinemia (MC, n = 10). Using HILIC-HRMS for polar metabolites and RP-UHPLC-HRMS for lipids, the authors identified distinctive metabolic and lipidomic signatures for HCC. Key findings included: Marked elevation of short- and long-chain acylcarnitines in HCC, consistent with altered mitochondrial β-oxidation; Pronounced reduction of lysophosphatidylcholines (LPCs), both saturated and unsaturated, suggesting dysregulation of phospholipid remodeling via the Lands’ cycle; Pathway enrichment highlighting mitochondrial β-oxidation of short-chain saturated fatty acids and phospholipid biosynthesis as significantly modulated in HCC; Supervised modeling (PLS-DA) demonstrating that combined metabolomics and lipidomics datasets outperform AFP for distinguishing HCC from HCV and MC, including AFP-negative cases, with AUC values up to 0.94.

Here, we re-analysed the publicly available lipidomics dataset from this study using the Functional Lipid Analysis (FLA) module of LipidOne.

### Functional alterations in HCC versus HCV chronic infection

To obtain an overview of functional lipid differences between HCC and chronic HCV infection, we inspected the bubble heatmap of the most perturbed FLA indices (Figure 4). Color encodes the log₂ fold change for HCV relative to HCC (red = higher in HCV; blue = lower), and bubble size scales with statistical support (−log₁₀ p-value). Each of the 10 main indices is accompanied by a concise sentence that facilitates its functional interpretation.

**Figure 4:**
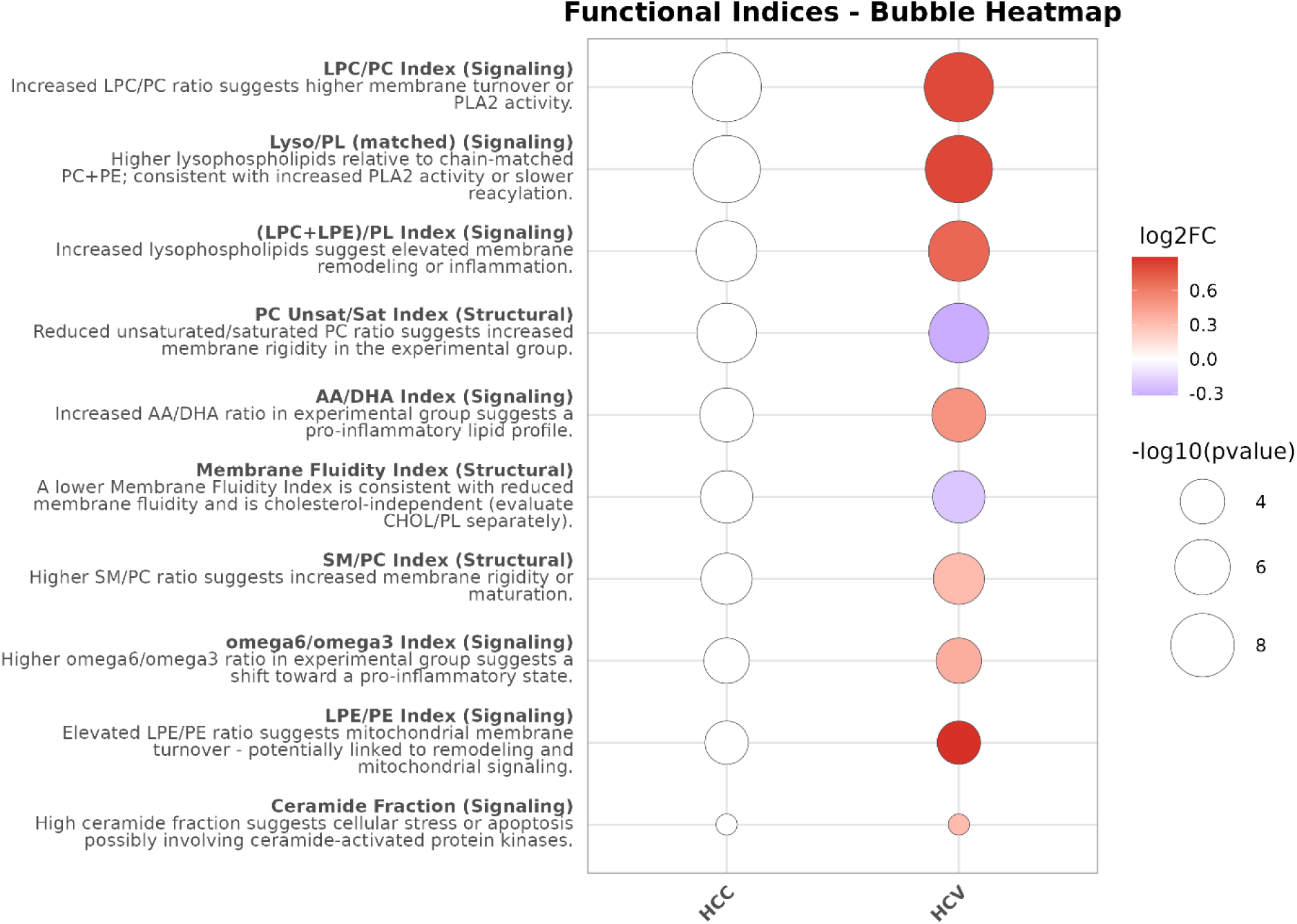
Functional lipid indices distinguishing chronic HCV infection from HCC. Bubble heatmap of FLA indices comparing HCV vs HCC. Bubble color shows log₂(HCV/HCC) (red = higher in HCV; blue = lower), and size is proportional to −log₁₀(p-value). Indices displayed: LPC/PC, Lyso/PL (matched), (LPC+LPE)/PL, PC Unsat/Sat, AA/DHA, Membrane Fluidity Index, SM/PC, omega-6/omega-3, LPE/PE, and Ceramide Fraction. HCV shows strong increases in lysophospholipid-based indices and in LPE/PE, consistent with elevated PLA2-mediated turnover and mitochondrial/inner-membrane remodeling. Decreases in PC Unsat/Sat and in the Membrane Fluidity Index indicate more rigid, less fluid membranes relative to HCC; SM/PC is higher in HCV, further supporting increased membrane order. AA/DHA and omega-6/omega-3 are elevated in HCV, supporting a pro-inflammatory shift. The Ceramide Fraction is slightly lower in HCV.

A clear predominance of signaling-related alterations emerged. HCV showed robust increases in multiple lysophospholipid indices—LPC/PC, (LPC+LPE)/PL, and Lyso/PL (matched)—consistent with enhanced PLA2 activity and/or slower reacylation compared with HCC. The LPE/PE index was also markedly elevated in HCV, suggesting increased turnover of inner/mitochondrial membranes.

In contrast, PC Unsat/Sat and the Membrane Fluidity Index decreased in HCV, indicating reduced membrane fluidity (greater rigidity); SM/PC was modestly higher in HCV, pointing in the same direction of increased membrane order. Pro-inflammatory balance indices (AA/DHA and omega-6/omega-3) were higher in HCV, consistent with a more pro-inflammatory lipid milieu. The Ceramide Fraction was slightly lower in HCV, arguing against enhanced ceramide-driven stress relative to HCC. Altogether, these patterns recapitulate the lysophospholipid depletion reported in HCC by Caponigro et al., while translating lipid changes into a functional framework linking membrane remodeling, inflammation, and organelle turnover.

### Identification of the most discriminant functional indices between HCC and HCV

The functional volcano plot (Figure 5) summarizes effect size [log2(HCV/HCC)] versus significance (−log10 p) across Energy, Signaling, and Structural indices. Consistent with Caponigro et al., we confirm a marked depletion of lysophospholipids in HCC: indices that rise in HCV include LPC/PC, (LPC+LPE)/PL, and Lyso/PL (matched), all exceeding both effect-size and significance thresholds. FLA adds two functional refinements not apparent from class-level lipid lists alone. First, the ether-specific Lyso-O/PL-O (matched) index is also higher in HCV, indicating that PLA2-linked turnover extends to ether lipids, a detail that implicates peroxisomal/ER pathways of membrane remodeling. Second, LPE/PE is strongly increased in HCV, pointing to heightened inner membrane/mitochondrial remodeling relative to HCC.

**Figure 5:**
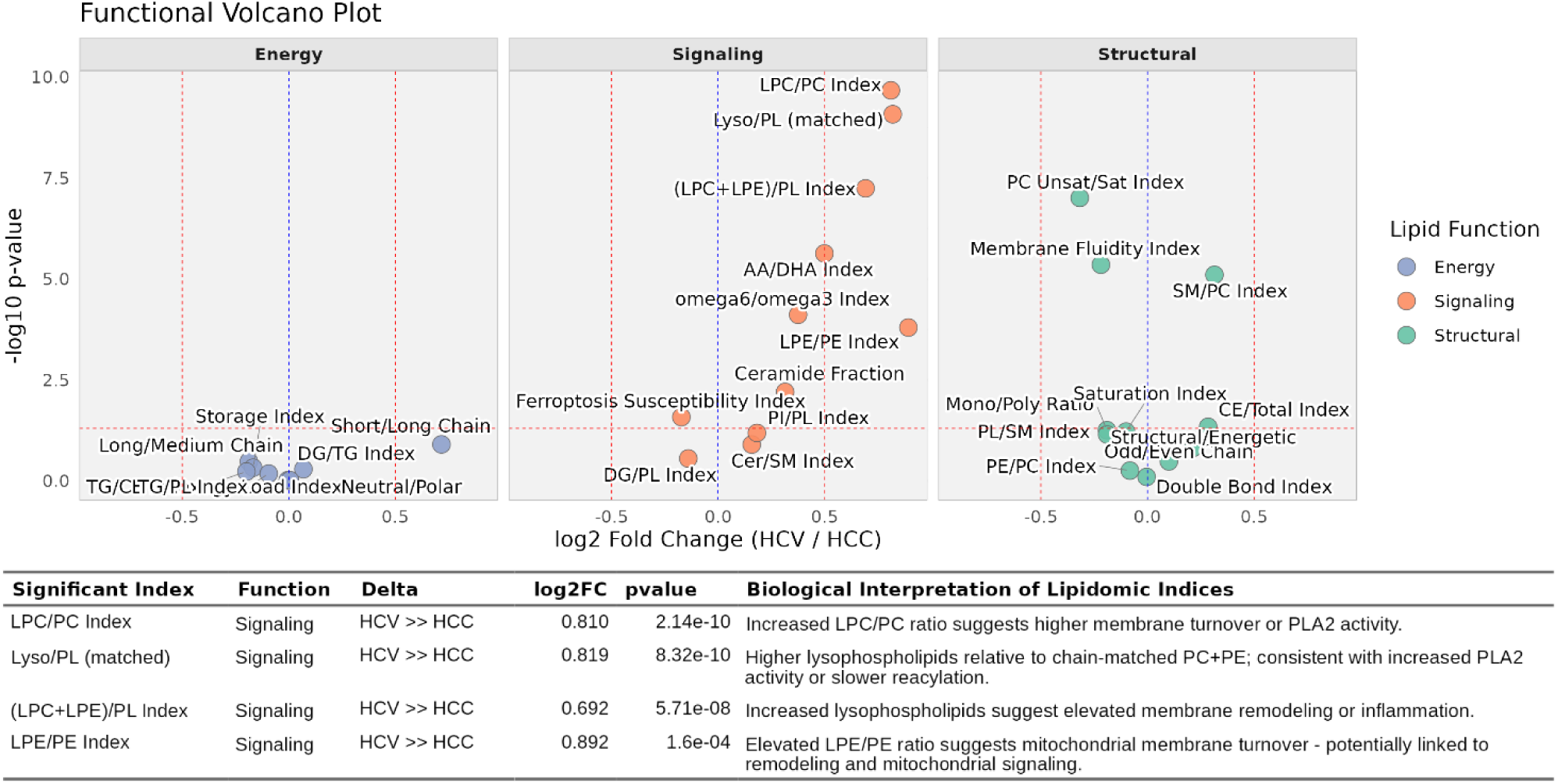
Functional Volcano Plot (HCV vs HCC). Each point is a Functional Lipid Analysis (FLA) index plotted by log₂(HCV/HCC) on the x-axis and −log₁₀(p-value) on the y-axis. Facets group indices by lipid function (Energy, Signaling, Structural). Vertical dashed lines mark the user-defined fold-change band; the horizontal dashed line marks p = 0.05. Labels identify indices that exceed both thresholds; the table below reports these hits with direction and statistics. Significant increases in HCV include LPC/PC, (LPC+LPE)/PL, Lyso/PL (matched), and LPE/PE, consistent with enhanced PLA2-linked membrane turnover and mitochondrial/inner-membrane dynamics. PC Unsat/Sat and the Membrane Fluidity Index trend lower in HCV, while SM/PC trends higher; AA/DHA and omega-6/omega-3 show pro-inflammatory shifts but remain within the effect-size band. The Ceramide Fraction shows only a small, sub-threshold difference.

Among signaling ratios, HCV displays four large-effect, significant increases—LPC/PC, (LPC+LPE)/PL, Lyso/PL (matched), and LPE/PE—supporting elevated PLA2 activity and/or slower reacylation together with increased turnover of inner/mitochondrial membranes. Pro-inflammatory balance indices (AA/DHA, omega-6/omega-3) tend to be higher in HCV but fall inside the preset effect-size band. The Ceramide Fraction shows only a modest, non-significant shift, providing no evidence for stronger ceramide-driven signaling in either group.

Structural trends include lower PC Unsat/Sat and a lower Membrane Fluidity Index, together with slightly higher SM/PC in HCV—patterns consistent with more ordered (less fluid) membranes—yet these remain below the combined thresholds. Energy-related indices cluster near the origin and do not pass the filters, indicating that the dominant differences between HCV and HCC are signaling/structural rather than storage-centric. Overall, the volcano plot corroborates the depletion of lysophospholipids in HCC reported by Caponigro et al. and reframes the lipid changes in a functional context that links membrane remodeling, inflammation, and organelle turnover.

### Functional biomarker discovery for HCC detection in HCV-positive patients

From the complete panel of Functional Lipid Analysis (FLA) indices, we selected the top 20 discriminators for HCV vs HCC (Figure 6). For each index we report p-value, ROC AUC, statistical power, and Cohen’s d to quantify diagnostic potential.

**Figure 6:**
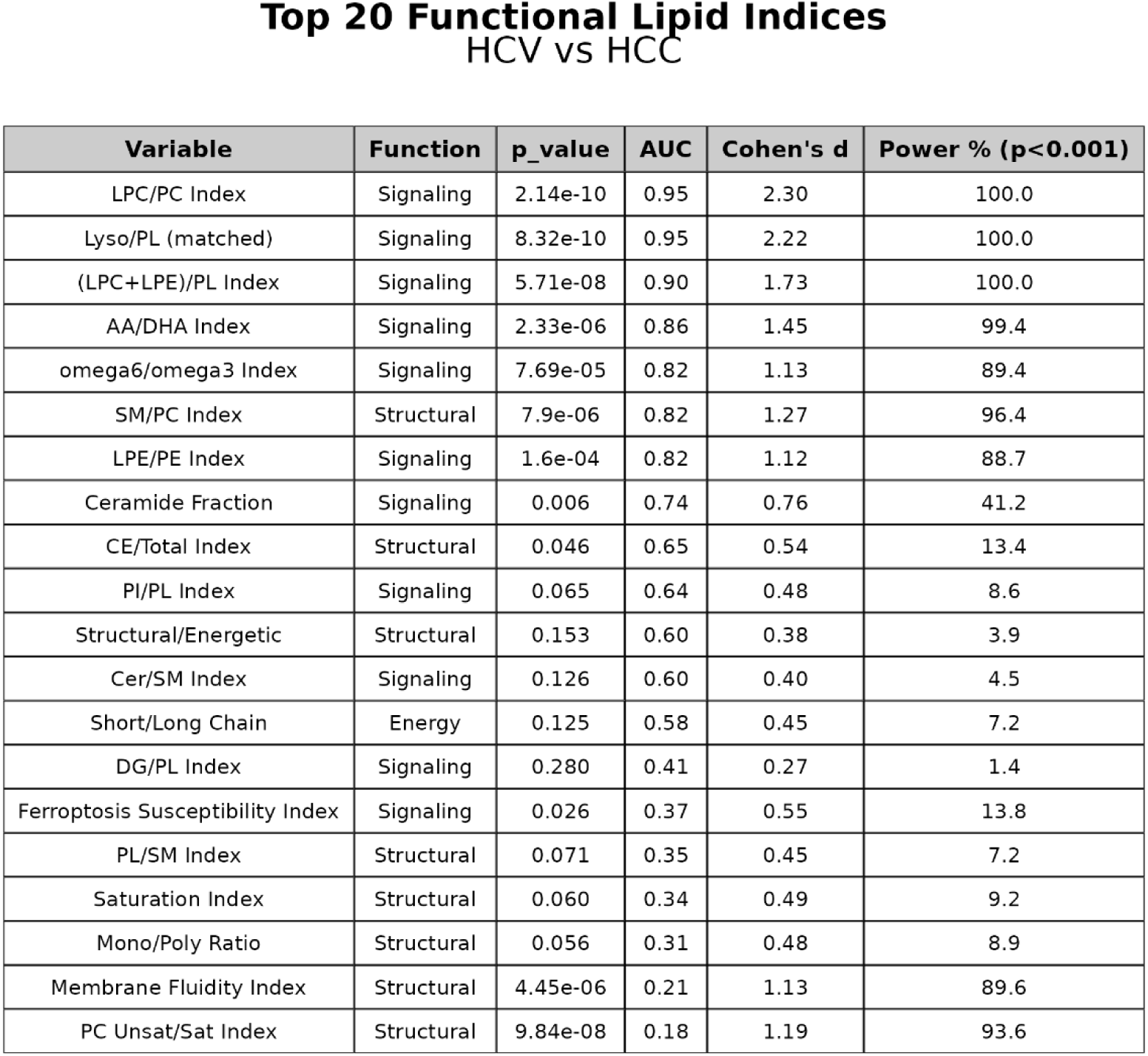
Top 20 functional lipid indices discriminating HCV from HCC. Table of the 20 highest-ranking FLA indices with Function, p-value, ROC AUC, Cohen’s d, and Power % (for p<0.001). Leading markers include LPC/PC and Lyso/PL (matched) (AUC ≈ 0.95; d > 2), followed by (LPC+LPE)/PL (AUC ≈ 0.90) and LPE/PE (AUC ≈ 0.82). Inflammatory ratios AA/DHA and omega-6/omega-3 also perform well (AUC ∼0.82–0.86). Structural readouts show strong inverse discrimination: PC Unsat/Sat and the Membrane Fluidity Index have AUC < 0.5 (≈0.18–0.21), consistent with lower unsaturation/fluidity in HCV relative to HCC, while SM/PC (AUC ≈ 0.82) is higher in HCV, indicating greater membrane order. Note: AUC < 0.5 indicates inverse polarity (marker higher in HCC); flipping the decision rule yields the complementary AUC (1 − AUC) and the same discriminative strength.

Consistent with Caponigro et al., signaling indices dominate the ranking. The strongest performers are LPC/PC and Lyso/PL (matched) (AUC ≈ 0.95; d ≥ 2.2), followed by (LPC+LPE)/PL and LPE/PE (AUC ≈ 0.90 and ≈ 0.82, respectively). Together these increases indicate elevated PLA2-driven membrane turnover and enhanced inner/mitochondrial-membrane dynamics in HCV. Inflammatory balance indices (AA/DHA, omega-6/omega-3) also rank highly, consistent with a more pro-inflammatory lipid milieu in HCV.

Structural readouts point to more ordered/less fluid membranes in HCV: PC Unsat/Sat and the Membrane Fluidity Index show strong effects with inverse polarity (AUC < 0.5), while SM/PC is higher in HCV and directionally concordant. The Ceramide Fraction shows moderate discrimination (p = 0.006; AUC ≈ 0.74) but does not exceed the preset effect-size threshold in the volcano plot. Energy-related indices cluster near the origin and do not pass the combined filters, indicating that the dominant HCV–HCC differences are signaling/structural rather than storage-centric.

Moreover, when applying PCA to the molecular species to explain the variations (Figure S2), the first principal component accounts for 20.3% of the variance, compared to 35.9% when computed on indices. These results once again confirm that indices enhance the analytical power and provide more interpretable information.

Overall, functional indices dampen single-species noise while retaining statistical power and mechanistic interpretability. This re-analysis confirms the lysophospholipid depletion in HCC reported by Caponigro et al. and reframes it within a functional taxonomy linking membrane remodeling, inflammation, and organelle turnover, yielding robust, clinically relevant candidates for HCC detection in HCV-positive populations.

## 4. Conclusion

Functional Lipid Analysis (FLA) turns untargeted lipidomes into mechanism-aware, organelle-aware readouts. Rather than listing hundreds of species, FLA quantifies a curated set of functional indices spanning signaling, structural, and energy domains and ties each index to a concise biochemical interpretation. The result is an analysis that tells not only what changed, but why it matters. FLA is not an enrichment analysis; it is mechanism-grounded inference from quantitative lipid features.

Across two independent case studies, FLA both replicated the key findings of the original works and extended them with insights that are difficult to obtain from class-level summaries alone. In cardiomyocytes, FLA separated storage-positive from mobilization-negative indices and mapped the phenotype onto a Pdk1–Pdk4– Pdha1 network interfacing with Pnpla2 (ATGL) and FAO enzymes. In HCV/HCC plasma, FLA confirmed lysophospholipid depletion in HCC and newly highlighted ether-lysophospholipid turnover, mitochondrial/inner-membrane remodeling (LPE/PE), and ceramide-axis differences. Because indices are ratio-based and statistics are attached to each readout (effect size, AUC, power), the outputs are interpretable, portable, and biomarker-ready.

Importantly, FLA binds statistical outputs to standardized mechanistic statements, converting lipidomic ratios into actionable interpretations that are accessible to clinical and translational teams without deep lipid biochemistry expertise.

To our knowledge, FLA is the first general-purpose bioinformatics module that (I) organizes lipidomes into functional axes, (II) overlays results on protein interaction networks to propose enzyme-level hypotheses, and (III) delivers a compact suite of publication-grade visuals (summary bars, bubble heatmap, functional volcano, top-index table, protein network) in minutes. This positions lipidomics not only as a descriptive assay but as a strategic tool for mechanism discovery and translational biomarker development.

## 5. Limitations and future directions

Accurate computation of several indices requires molecular-species–level annotation (chain length/unsaturation and, where relevant, O-/P-linkages). Datasets reported only as sum compositions limit index coverage.

Index accuracy depends on class coverage and identification quality. Ratio-based design mitigates but does not eliminate variability due to extraction, chromatography, adduct handling, or missing values (zero handling can inflate ratios at low abundance).

Some indices are correlated by construction; they should be interpreted as a panel rather than independent tests. Statistical control remains essential.

FLA infers function from correlations in lipidomes and maps results onto STRING; it does not measure enzyme activity. Network-level hypotheses (e.g., ATGL, SCD1, ELOVLs) require orthogonal validation (proteomics, flux assays).

Finally, the index library is curated and finite: rare contexts or novel lipid chemistries (e.g., oxidized species, cardiolipin microheterogeneity) may not yet be captured.

Future work will expand the index library (including oxidized lipids and pathway-specific composites), add data-driven index discovery and uncertainty quantification, tighten cross-study harmonization (QC rules, FDR defaults, zero-handling), and deepen multi-omics integration (proteome, transcriptome, 13C-flux) so that FLA can progress from functional readouts to testable, enzyme-level mechanisms at scale.

## Availability

New FLA module is freely available through the LipidOne.eu web platform.

## Author contributions

H.B.R.A.: Conceptualization, Methodology, Writing – original draft, Writing – review & editing.

D.F.M.: Validation, Data curation.

M.P.: Validation, Data curation.

L.M.: Writing – review & editing, Resources.

S.M.: Writing – review & editing.

A.K.: Writing – review & editing, Resources.

R.M.P.: Conceptualization, Methodology, Software, Visualization, Writing – original draft, Supervision.

## Conflicts of Interest

The authors declare no conflicts of interest.

## Funding

This research was supported by the Wellcome Trust [grant number 226800/Z/22/Z], the NIHR Cambridge Biomedical Research Centre (NIHR203312), and the HDM-FUN project (European Union’s Horizon 2020 research and innovation programme, grant number 847507). The views expressed are those of the authors and not necessarily those of the NIHR, the Department of Health and Social Care, or the European Union.

## Supporting information

Lipidomic Dataset 1

Lipidomic Dataset 2

## Acknowledgement

we would like to thank Matteo Boschi (UMMON.it) for his invaluable cooperation in developing the LipidOne 2.3 website front-end and coding its back-end interactions with the R scripts.

## Notes

### Competing Interest Statement

The authors have declared no competing interest.

