## Supplemental Figure S1 and S2 for "Functional Lipid Analysis via Index-Based Lipidomics Profile: A New Computational Module in LipidOne"

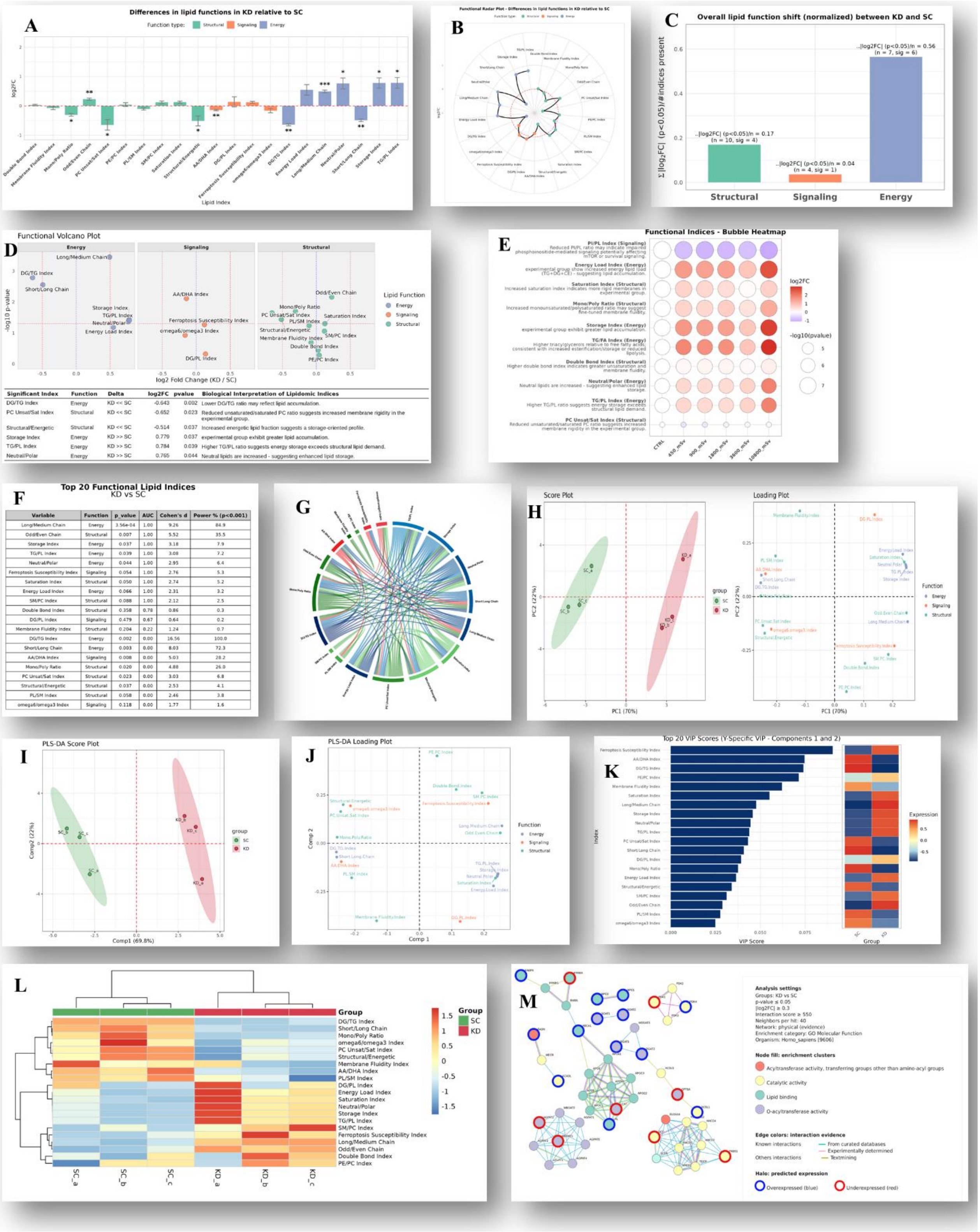


*Figure S1: Examples of graphical outputs generated by the FLA module in LipidOne. (A) Bar plot comparing all indices; (B) Radar plot summarizing the overall functional landscape; (C) Functional Dominance plot showing a three-bar panel (Structural, Signaling, Energy) to study functional shifts within a study; (D) Functional Volcano plot; (E) Functional Bubble plot displaying a bubble heatmap of top index differences across groups; (F) Functional biomarker discovery table; (G) Functional Correlation Circos plot; (H) PCA score and loading plots; (I) PLS-DA score plot; (J) PLS-DA loading plot; (K) VIP score bar plots; (L) Functional heatmap of indices; (M) Predicted functional protein network.*

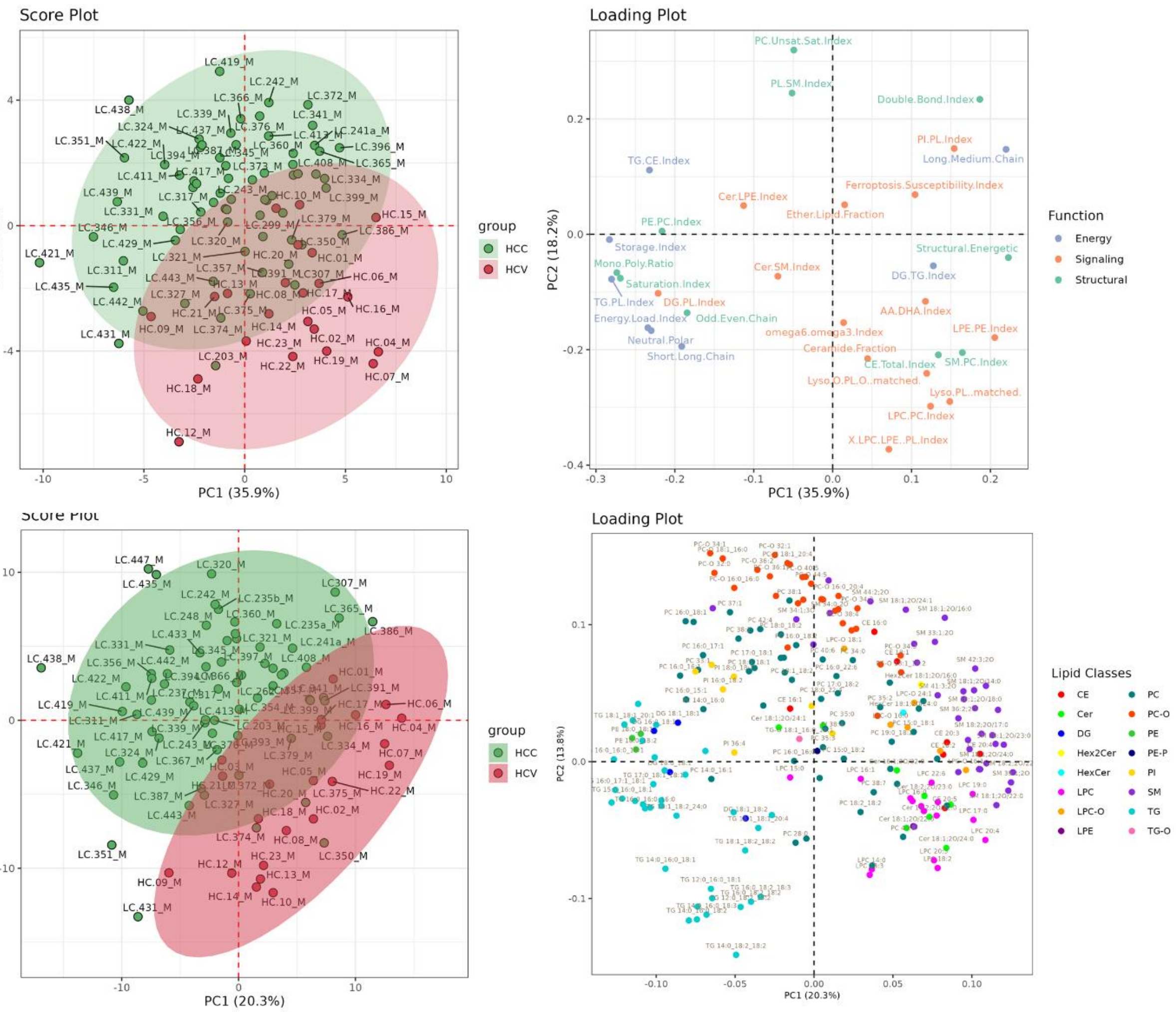


*Figure S2: PCA on functional indices versus molecular species. Top: PCA computed on the matrix of functional indices (score plot left; loading plot right). Bottom: PCA computed on molecular lipid species from the same samples (score left; loading right). Both analyses use identical preprocessing (centred and auto scaled). Score plots show 95% confidence ellipses for groups; axes report the percentage of explained variance. In this dataset, PC1 explains 35.9% of the variance with indices and 20.3% with molecular species, illustrating how index-based features sharpen group separation and yield more interpretable loadings (indices coloured by function; species coloured by lipid class).*
